# Zika virus capsid protein C (ZIKV-C) interactors network mapped by proteomic analysis of human neural stem cells expressing FLAG-tagged C protein

**DOI:** 10.1101/2025.07.24.666584

**Authors:** Agata Malinowska, Magdalena Bakun, Agnieszka Brzozowska, Ewa Liszewska, Malgorzata Podsiadla-Bialoskorska, Bianka Swiderska, Ewa Sitkiewicz, Ewa Szolajska, Michal Hetman, Michal Dadlez, Jacek Jaworski

**Author notes:** Corresponding authors: **Jacek Jaworski**, Laboratory of Molecular and Cellular Neurobiology, International Institute of Molecular and Cell Biology, Ks. Trojdena St. 4, Warsaw, 02-109, Poland, **Michał Dadlez**, Mass Spectrometry Laboratory, Institute of Biochemistry and Biophysics, Polish Academy of Sciences, 5A Pawinskiego St., 02-106, Warsaw, Poland.

## Abstract

Zika virus is a teratogenic pathogen belonging to the *Flaviviridae* family. It possesses the ability to penetrate the placenta and affect the brain development of a fetus, resulting in microcephaly and functional impairments. Mechanisms of this neurotoxicity are still unclear, but capsid proteins of Zika and related viruses are known to exert apoptotic effect in different types of cells, including neurons. To explore the pathways affected by the presence of ZIKV-C, we have performed MS-based interactomic experiment in human neural stem cells and managed to identify 149 putative interactors. Our results indicate that the nucleus (especially the nucleolus) and the mitochondria are the main sites of interaction of protein C with host proteins. A number of the proteins we identified have significant links to diseases of the nervous system, including neurodevelopmental diseases. Furthermore, for the first time, we have identified MAM-domain containing glycosylphosphatidylinositol anchor protein 1, T-complex protein 1 subunit beta, lysine-tRNA ligase, calumenin as particularly abundant ZIKV protein C interactors. Data are available via ProteomeXchange with identifier PXD064412.

## Introduction

Zika virus (ZIKV) is an RNA virus from the Flaviviridae family, associated with the microcephaly epidemic in Brazil during 2015–2016. Infections occurring during the first and early second trimesters of pregnancy (and, to a lesser extent, later stages) can lead to brain malformations. Birth defects—including cortical atrophy with microcephaly and functional impairments such as dysphagia and epilepsy—are reported in 5–15% of babies born to women infected during pregnancy^1^. This neurodevelopmental disruption is consistent with the neurotropic nature of other arthropod-borne flaviviruses related to ZIKV^2^. For instance, mosquito-transmitted West Nile virus (WNV) and Japanese encephalitis virus (JEV) infect neurons of the central nervous system, causing neurological diseases that may involve neuronal loss and lasting brain function deficits^3^. However, ZIKV appears to be a uniquely potent brain teratogen. It is known to penetrate the placenta, infect the fetal brain, and exhibit a strong capacity to infect neural stem cells (NSCs), neuroprogenitor cells, and developing neurons—impairing their growth and inducing apoptosis.

The ZIKV virion comprises an RNA genome and three structural proteins: capsid (C), membrane (M), and envelope (E). The ZIKV-C protein, the first flaviviral protein translated upon infection, forms the core of the virion by associating with viral RNA^8^. Beyond its structural role, ZIKV-C is believed to facilitate infection. It has been shown to suppress the expression of type I interferon and related genes^9^, and to form complexes with stress granule components such as Ras GTPase-activating protein-binding protein 1 and Caprin-1, thereby inhibiting stress granule formation^10^. ZIKV-C also binds to the nuclear Up-frameshift protein 1 (UPF1), a key regulator of nonsense-mediated mRNA decay^11^. While these interactions enhance viral survival in host cells, they are also likely to contribute to central nervous system pathology, including microcephaly—highlighting the need to better understand ZIKV-host protein interactions.

Although ZIKV replicates in invaginations of the endoplasmic reticulum (ER) membrane, some flaviviral proteins localize to the nucleus of infected cells, where they enhance virus production^12–14^. Nucleolar localization has been reported for the capsid protein in ZIKV^15^ and related viruses such as WNV and JEV^16^. Additionally, WNV- and JEV-C proteins interact with ribosome biogenesis factors like nucleophosmin, nucleolin, and the ATP-dependent RNA helicase DDX56^12–14^, contributing to viral replication. Nuclear localization of the flaviviral capsid protein has also been associated with cytotoxic effects. In WNV-infected SH-SY5Y neuroblastoma cells, nucleolar WNV-C was found to sequester and inhibit the p53 ubiquitin ligase Mdm2^18^, thereby activating the apoptotic p53 pathway. Both nucleolar localization and pro-apoptotic effects have similarly been observed for ZIKV-C^19^.

Several interactomes of ZIKV proteins, including that of ZIKV-C, have been published^20–22^, offering new insights into the regulation of the viral life cycle and the mechanisms underlying ZIKV-induced neurodevelopmental disruption. Yet the molecular mechanisms underlying ZIKV-induced neurodevelopmental disorders remain incompletely understood. Therefore, in this study, we used mass spectrometry to identify the ZIKV-C interactome in human NSCs. Several novel interactors were identified, including NSC proteins with known or potential roles in developmental neurogenesis.

## Materials and Methods

### Cell culture

The skin biopsy from a healthy volunteer was obtained with informed and written consent and processed anonymously. The study was approved by the Ethics Committee of the Children’s Memorial Health Institute, Warsaw, Poland (decision no. 112/KBE/2013). All experimental procedures/methods were performed in accordance with the relevant guidelines and regulations.

Induced pluripotent stem cells were obtained as previously described by Liszewska et al.^1^; in brief, human fibroblasts derived from skin biopsy (male, 40 years old) were cultured in Dulbecco’s modified Eagle’s medium (DMEM): high glucose medium supplemented with 10% fetal bovine serum, 1% penicillin-streptomycin (Sigma-Aldrich). The culture was maintained at 37°C in a humidified atmosphere containing 5% CO_2_. After the fourth cell passage, the fibroblasts were split and seeded at a density of 1 × 10^4^ cells/cm^2^. After 24 h, the cells were transduced in the presence of 5 µg/mL Polybrene (Sigma-Aldrich) with a single lentiviral vector expressing four transcription factors (Octamer-binding transcription factor 4, Krüppel-like factor 4, Transcription factor SOX-2 and c-Myc) from a single transcript for the next two days. At 48 h post transduction, the fibroblasts were plated on a feeder layer of mouse embryonic fibroblasts (MEFs) (EmbryoMax Primary Mouse Embryonic Fibroblasts-PMEF-CFL, Sigma-Aldrich) inactivated with mitomycin C (Sigma-Aldrich) and cultured in iPSC medium (DMEM/F12, 20% knockout serum replacement, 1% non-essential amino acid stock, 1 mM GlutaMax, 1 mM sodium pyruvate, 100 µM β-mercaptoethanol, 1% penicillin-streptomycin (Thermo Fisher Scientific) and 10 ng/mL basic fibroblast growth factor (bFGF, Alomone Labs). The iPSC medium was changed every other day for the next 3 weeks. After 21 days, iPSC colonies were manually collected and seeded onto Matrigel-coated plates (Corning) in Essential 8 medium (Thermo Fisher Scientific) with 10 µM ROCK inhibitor (Tocris Bioscience) for 24 hours. After this time, iPSCs were passaged every 4-5 days with Versene solution (Thermo Fisher Scientific) in the presence of the ROCK inhibitor for 24 hours and cultured in Essential 8 medium.

The iPSCs were differentiated into NSCs using the embryoid body [EB]-based protocol. To generate EBs, the iPSC colonies were detached using Versene solution, washed with phosphate-buffered saline, transferred to a low-attachment culture dish (Corning) and cultured in suspension in iPSC medium without bFGF for five days, with half of the medium changed every other day. The EBs were then attached to Matrigel coated plates and cultured in NSC medium consisting of DMEM/F12, N2 supplement (1:100), B27 supplement (1:100), 1 mM GlutaMax, 1% penicillin-streptomycin (Thermo Fisher Scientific), 20 ng/mL bFGF and 20 ng/mL epidermal growth factor (EGF, Alomone Labs). After five days, the emerging neural rosette structures were manually isolated, dispersed by pipetting and plated on Matrigel coated plates in NSC medium. After reaching confluence, NSC cultures were passaged with Accutase (Sigma-Aldrich). Cells were maintained in NSC medium, on Matrigel coated plates, at 37°C in a humidified atmosphere with 5% CO_2_ and passaged at 70-80% confluence using Accutase enzyme in the presence of 10 µM ROCK inhibitor for 24 hours.

### Plasmids

Two plasmids were used for the transfection of cultured NSCs: pLV_Zika_Cv_Flag was a gift from Vaithi Arumugaswami (Addgene) and pEGFP-C1 (Clontech).

### Transfection

For plasmid DNA transfection NSCs were seeded at 2×10^6^ per 100 Ø mm plate (VWR), which allowed approximately 60-80 % confluence to be reached within 24 hours. The following day, the cells were transfected with the pLV_Zika_Cv_Flag plasmid encoding Zika virus protein C (Zika virus, Asian genotype, PRVABC59 strain) with the Flag tag fused to the C-terminus or with the pEGFP-C1 control plasmid. The jetOPTIMUS transfection reagent (Polyplus) was used according to the manufacturer’s instructions with a plasmid to transfection reagent ratio of 1:1.5. To maximize transfection efficiency during the 4-hour incubation EGF, bFGF and antibiotics were not added to the transfection medium and the concentration of the N2 and B27 supplements was reduced to 0.1%. Afterward the medium was changed to standard NSC medium.

### Immunoprecipitation

At 48 hours after transfection NSCs were washed twice with DMEM/F12 and once with Tris-buffered saline (TBS: 25 mM Tris-HCl, pH 7.6; 100 mM NaCl). The cells were than collected by scraping in pre-cooled RIPA buffer (50 mM Tris-HCl, pH 7.6; 150 mM NaCl; 1 mM EDTA; 1% NP40; 5% glycerol) supplemented with cOmplete™EDTA-free Protease Inhibitor Cocktail (Roche) and were lysed at 4°C for 10 min on a rotary shaker, followed by centrifugation at 14,000 rpm for 10 min. Cleared lysates from transfected cells growing on two 100 Ø mm plates were mixed with 20 µl of Pierce™ Anti-DYKDDDDK Magnetic Agarose (Thermo Fisher Scientific) and incubated for 2 hours at room temperature on a rotary shaker. The beads with bound proteins were washed three times for 10 minutes at room temperature, with 100 µl of wash buffer (50 mM Tris-HCl, pH 7.6; 300 mM NaCl; 1 mM EDTA; 0.1% NP40). The FLAG-ZIKV-C protein and its interactors were then eluted using low-pH elution buffer, which is a component of the Pierce Crosslink Magnetic IP/Co-IP Kit (Thermo Fisher Scientific). For maximum recovery two 5-minute incubations at room temperature with 100 µl of elution buffer with constant rotation were performed. A magnetic stand was used throughout the procedure to separate the beads from the solution.

### Sample preparation for mass spectrometry analysis

Protein eluates were processed using the single-pot solid-phase-enhanced sample preparation (SP3) method with some modifications^2^. Cysteines were reduced by 1 hour incubation with 10 mM tris(2-carboxyethyl)phosphine (TCEP) at 60°C, followed by 10 min incubation at room temperature with 30 mM methyl methanethiosulfonate (MMTS). The magnetic beads mix was prepared by combining equal parts of Sera-Mag Carboxyl hydrophilic and hydrophobic particles (Cytiva). The beads mix was washed three times with MS-grade water and resuspended to a working concentration of 10 µg/µl. Twenty-five microliters of the prepared bead mix were added to each sample, along with phase B (0.1% formic acid in acetonitrile) to a final concentration of 80%. Proteins bound to beads were washed three times with 80% ethanol and once with acetonitrile. Dried resin was suspended in 50 µl of 100 mM ammonium bicarbonate buffer with 1 µg of trypsin (Promega). The first phase of the digestion was performed overnight at 37°C. For the second digestion cycle, an additional 0.5 µg of trypsin was added to the samples and the reaction was performed for additional 14 hours. After digestion, peptide solutions were transferred to new tubes, and the resin was further washed two times with 40 µl 1% DMSO in water and 60 µl water. Pooled peptide eluates were acidified to a concentration of 0.2% formic acid.

### Mass spectrometry

Peptides were analyzed using an LC-MS system comprising an Evosep One (Evosep Biosystems, Odense, Denmark) LC coupled to an Orbitrap Exploris 480 mass spectrometer (Thermo Fisher Scientific). Each sample was processed by loading half onto disposable Evotip Pure C18 trap columns (Evosep Biosystems) following the manufacturer’s protocol. The peptides bound to the columns were washed three times with 100 µl and subsequently covered with 300 µl of solvent A (0.1% formic acid in water). Chromatographic separation was carried out using an preformed 88-minute gradient (15 samples per day) on an analytical column (ReproSil Saphir C18, 1.5 µm beads, 150 µm ID, 15 cm length, Bruker Daltonics) at a flow rate of 220 nl/min. Data acquisition was performed in a positive mode with a data-dependent method employing following parameters: MS1 scans were acquired with a resolution of 60 000 and a normalized AGC target set to 300%, with an auto maximum inject time and a scanning a range from 300 to 1600 m/z. MS2 scans were performed with a resolution of 15 000, using a standard normalized AGC target and auto maximum inject time. The top 40 precursor ions within an isolation window of 1.6 m/z were selected for MS/MS analysis. Dynamic exclusion was set to 20 seconds with a mass tolerance of ±10 ppm, and a precursor intensity threshold of 5000 was applied. Fragmentation was performed in HCD mode with a normalized collision energy of 30%. Source settings included a spray voltage of 2.1 kV, a funnel RF level of 40, and a heated capillary temperature of 275 °C.

### Data analysis

Protein identification and quantification were performed with MaxQuant/Andromeda suite^**3**^ (version 2.5) using human reference protein database from Uniprot (proteome ID: UP000005640, version 2024_03, one sequence per gene, 20,590 sequences) supplemented with Zika virus proteins (Q32ZE1, Polyprotein divided into 14 sequences) and contaminants database. Reversed decoy database was used for false discovery rate (FDR) computation. LFQ quantification was allowed with classic normalization and “Match between runs” option was disabled. Other parameters were as follows: enzyme – Trypsin/P, static modifications – Methylthio (C), variable modifications – Oxidation (M), individual peptide tolerance – enabled, PSM and protein FDR – 0.01, min unique peptides - 1.

Further data analysis was performed in Perseus^4^ (version 1.6.15). Proteins identified only by modification site, reverse proteins and contaminants were removed. Proteins were log2 transformed and filtered to have 70% data completeness as a preparation for missing values replacement from normal distribution (separately for each column, width 0.3, down shift 1.8). Batch effect created by separate passages was removed with Limma algorithm, intensities were normalized to median. Two-sample Student’s test with permutative FDR was performed to identify proteins enriched in FLAG group. Proteins with q-value below 0.05 and difference above 0.49 (1.4-fold change) were submitted to upstream analysis with STRING (version 12.0) and Metascape ^5^ (version 3.5.20240101).

## Results

Six samples from NSCs expressing FLAG-ZIKV-C protein were analyzed against 6 control samples to search for potential interactors of viral capsid protein. NSC culture was repeated in parallel in 6 biological replicates of FLAG-ZIKV-C cells, paired with GFP controls. ZIKV-C protein was absent in controls and represented by 5-6 peptides (8-14 MS/MS spectra) in bait samples, attesting to robust expression in NSC samples.

Seven hundred sixty-seven human proteins were identified with 70% quantitative data completeness and at least 1 unique peptide (Supplement 1). T-test with permutative FDR was performed and yielded 350 significant (q value below 0.05) proteins, of which 165 were enriched in FLAG samples (Figure 1). Among these, 149 proteins showed enrichment greater than 40%. This dataset was subsequently used for upstream analyses using Gene Ontology (GO) tools and protein–protein interaction databases. The full quantification results are provided in Supplement 1.

**Figure 1:**
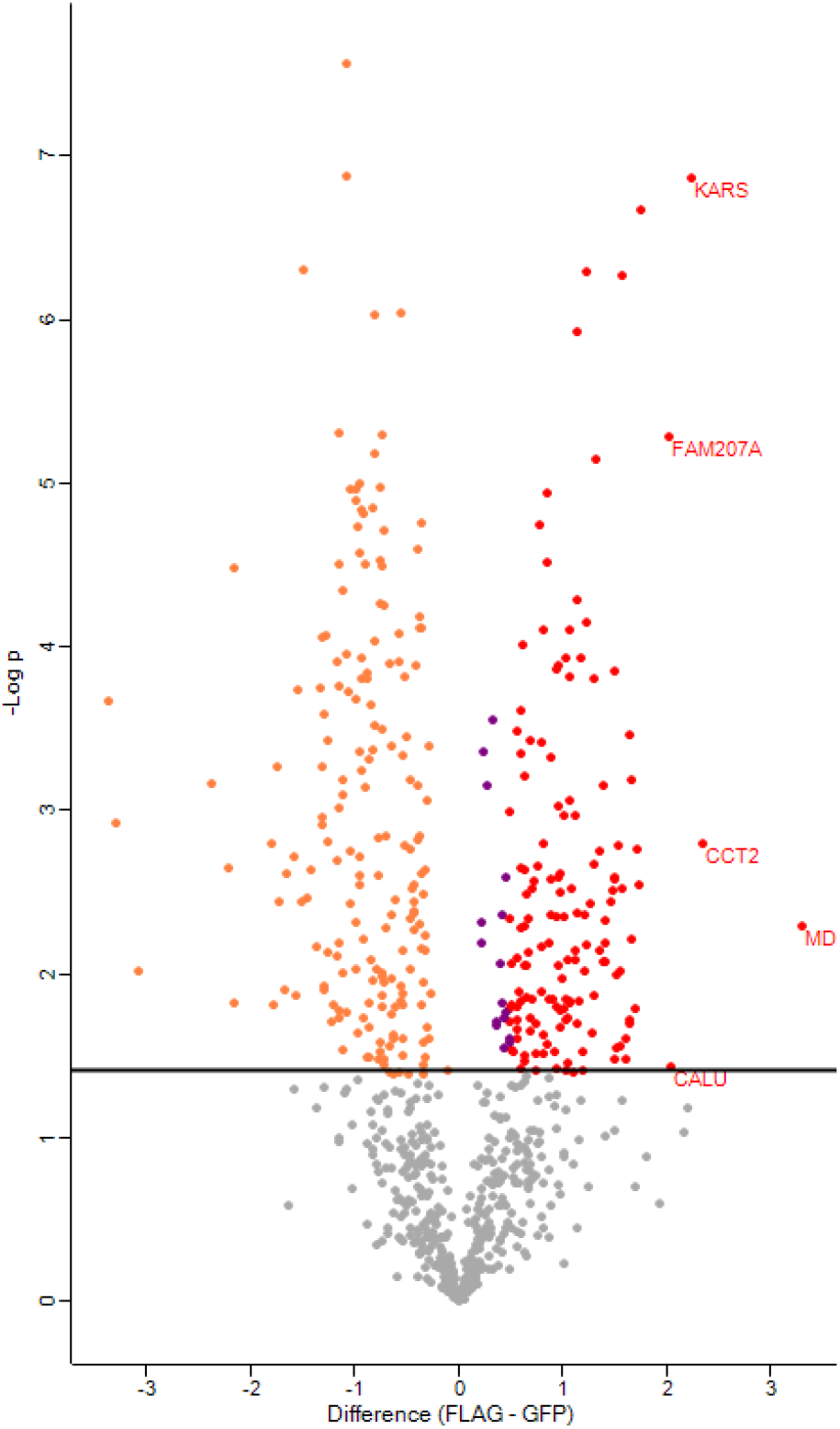
Volcano plot presenting quantitative differences between FLAG-ZIKV-C protein and controls. Log2 protein ratio is presented at X axis, −log10 p-value – at Y axis. Horizontal line denotes significance. Yellow dots represent proteins with decreased abundance, violet and red dots enriched proteins. Proteins with ratio exceeding 1.4 are marked red. Top 5-fold changes are labelled with Uniprot ID.

Enriched proteins with fold increase greater than 1.4, considered putative ZIKV-C interactors, were taken for further analysis using interactomic database STRING and Metascape software. A significant part of analysed proteins was assigned nuclear (96) or nucleolar (24) localization. Thirty one genes carried the GO Cell Component term “Mitochondrial” (Figure 2). Gene expression and translation-related terms dominate the GO Biological Process analysis (Figure 3), with significant representation of ribosomal proteins (mainly small ribosomal subunit components), part of them mitochondrial (mainly large subunit components).

**Figure 2:**
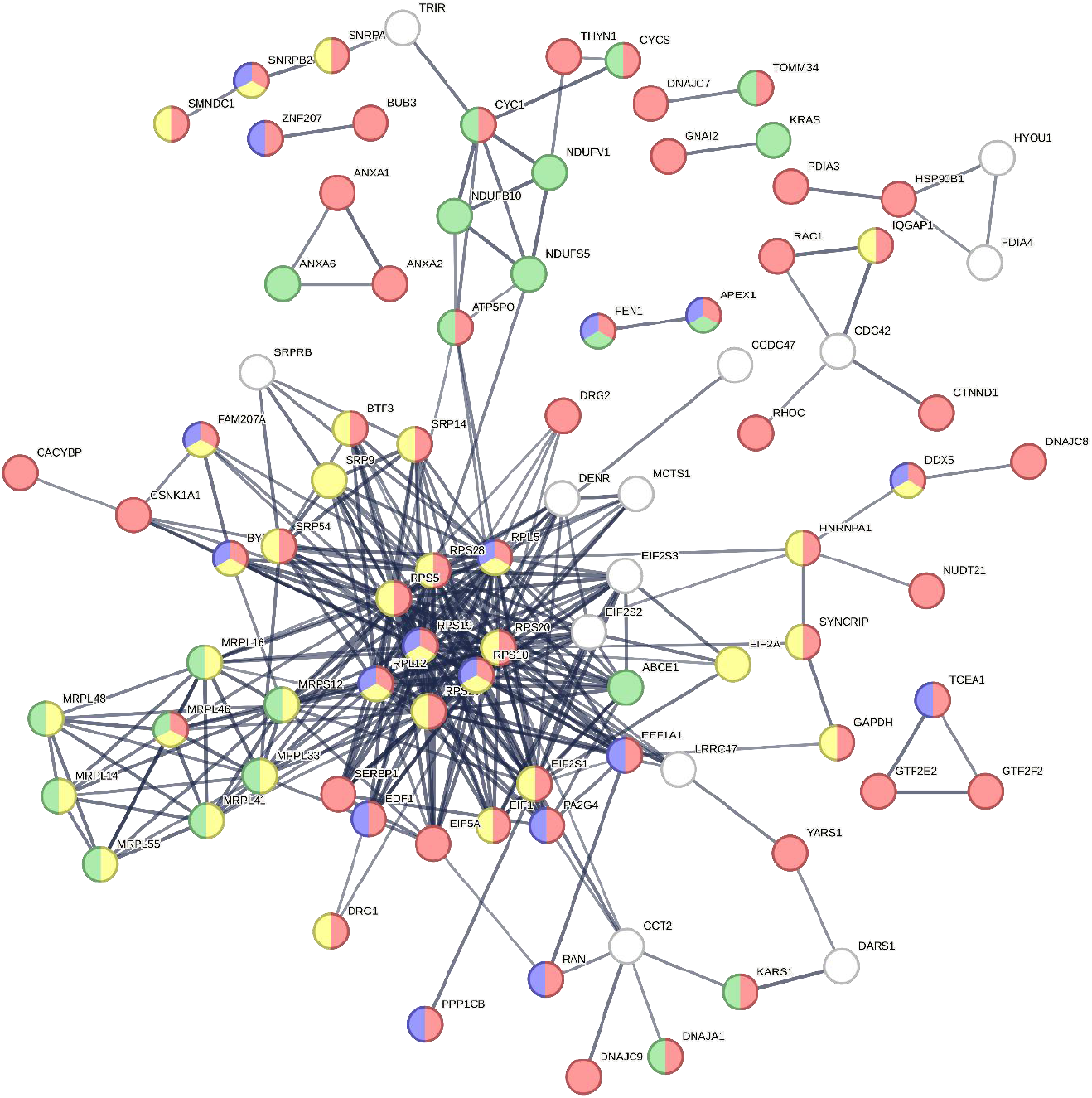
STRING results with GO Cellular Component visualization. 149 gene IDs were supplied to STRING, with following parameters: allowed data sources – Experiments, Databases, Co-expression, required confidence – high (0.7), hide disconnected nodes. Most proteins in the graph (96) are associated with nucleus (red, GO:0005634), with 24 qualified as nucleolar (blue, GO:0005730). Prominent cluster in the center comprises mostly of ribonucleoprotein complex (yellow, GO:1990904), with small mitochondrial subnetwork (green, GO:0005739).

**Figure 3:**
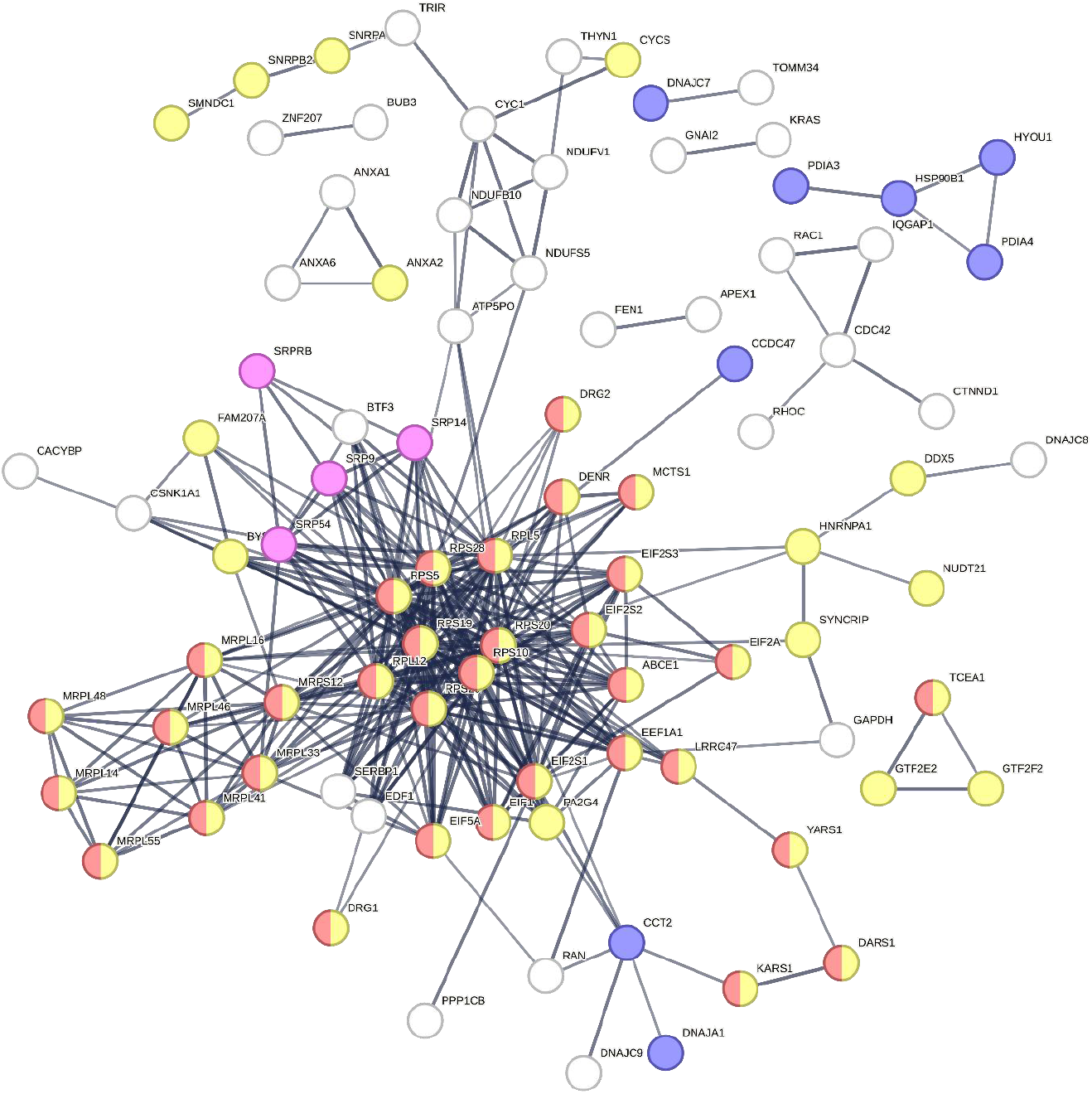
STRING results with GO Biological Process visualization. 149 gene IDs were supplied to STRING, with following parameters: allowed data sources – Experiments, Databases, Co-expression, required confidence – high (0.7), hide disconnected nodes. Significant part of putative ZIKV-C interactors is related to gene expression (yellow, GO:0010467), with marked presence of proteins involved in translation (red, GO:0006412), protein folding (blue, GO:0006457) and protein localization to endoplasmic reticulum (pink, GO:0072599).

Metascape analysis (Figure 4, Table 1) further emphasizes strong representation of translational machinery within the putative interactors dataset. Other important subgroups are associated with RNA processing/splicing, as well as mitochondrial ribosomal machinery and ATP synthesis.

**Table 1.**
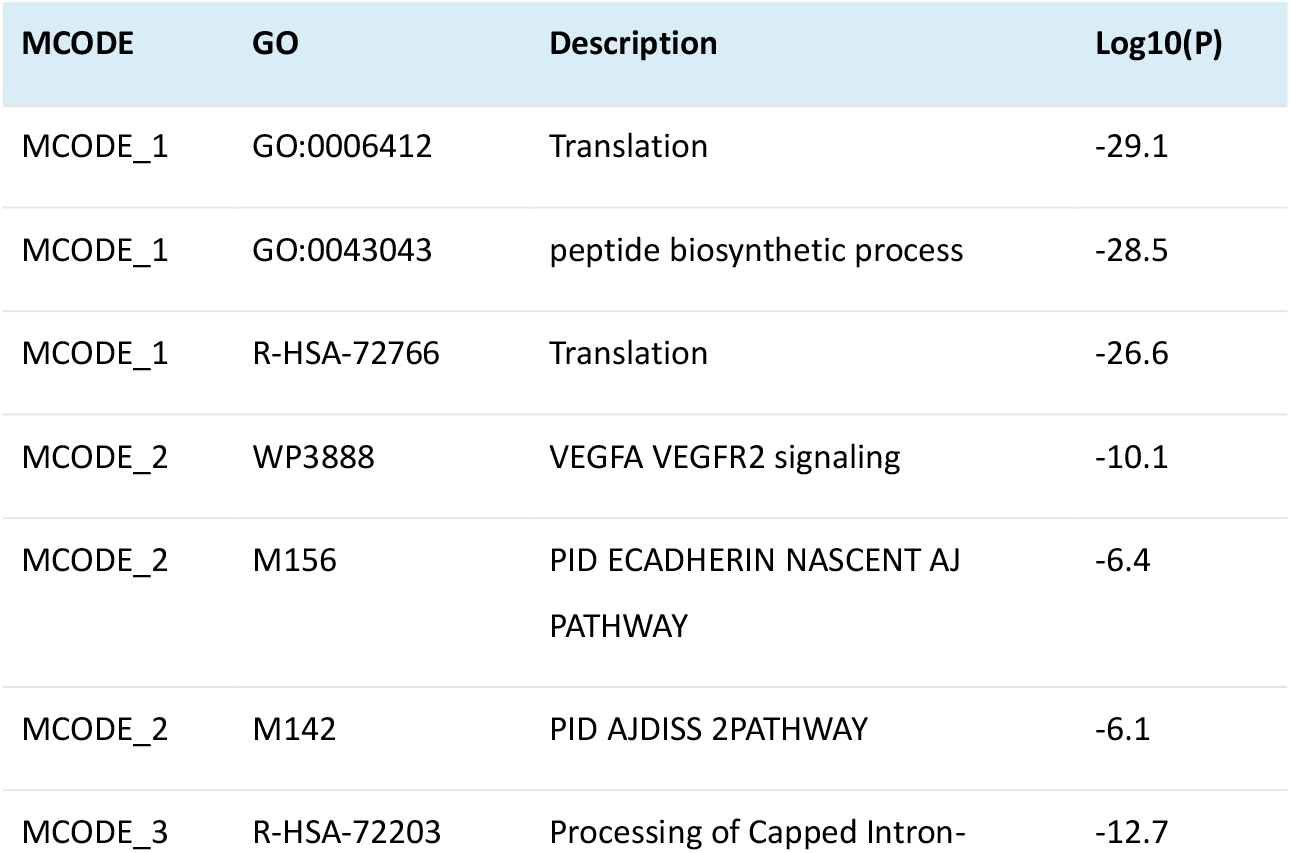
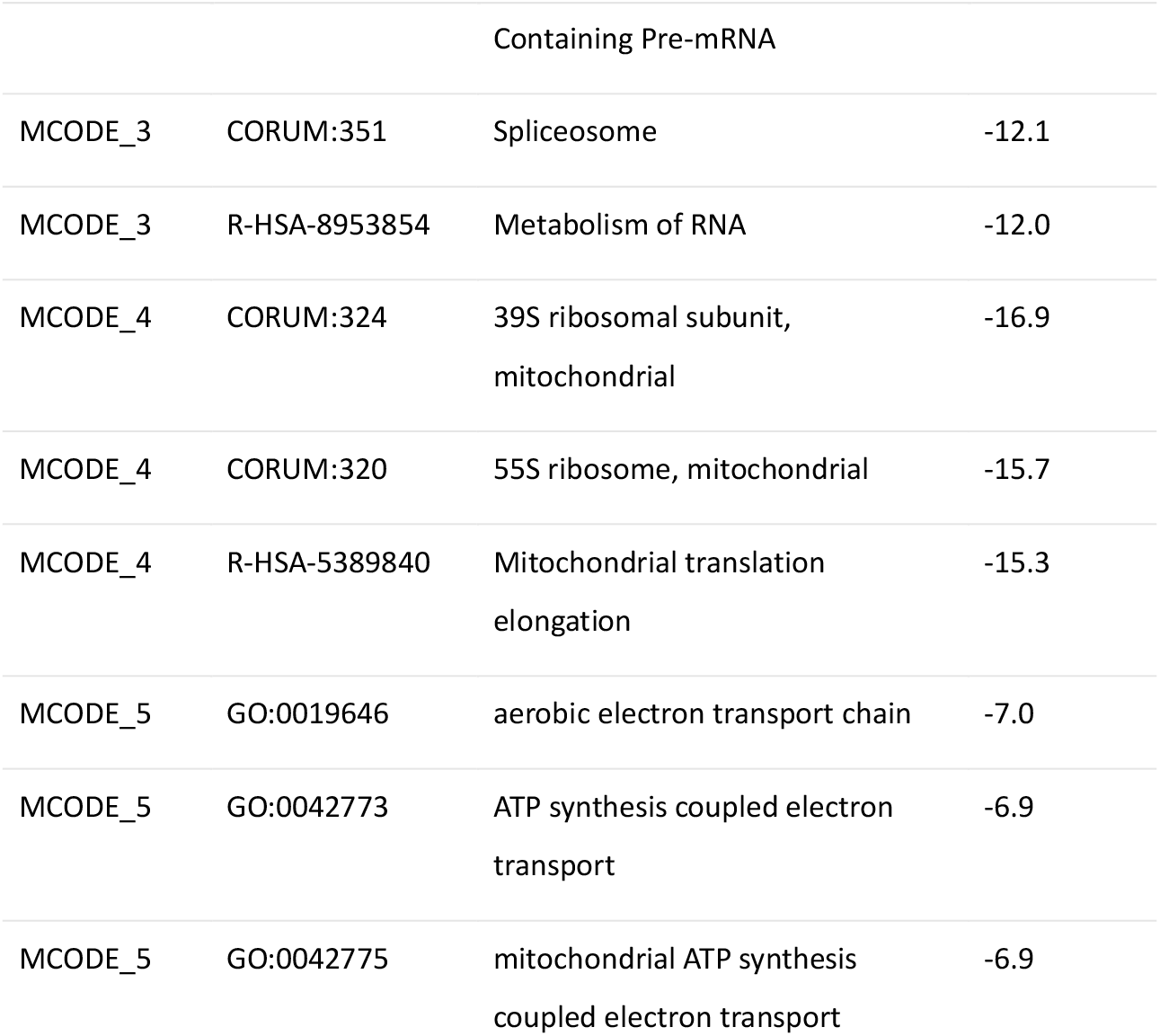
MCODE clusters and their respective terms in Metascape analysis.

**Figure 4:**
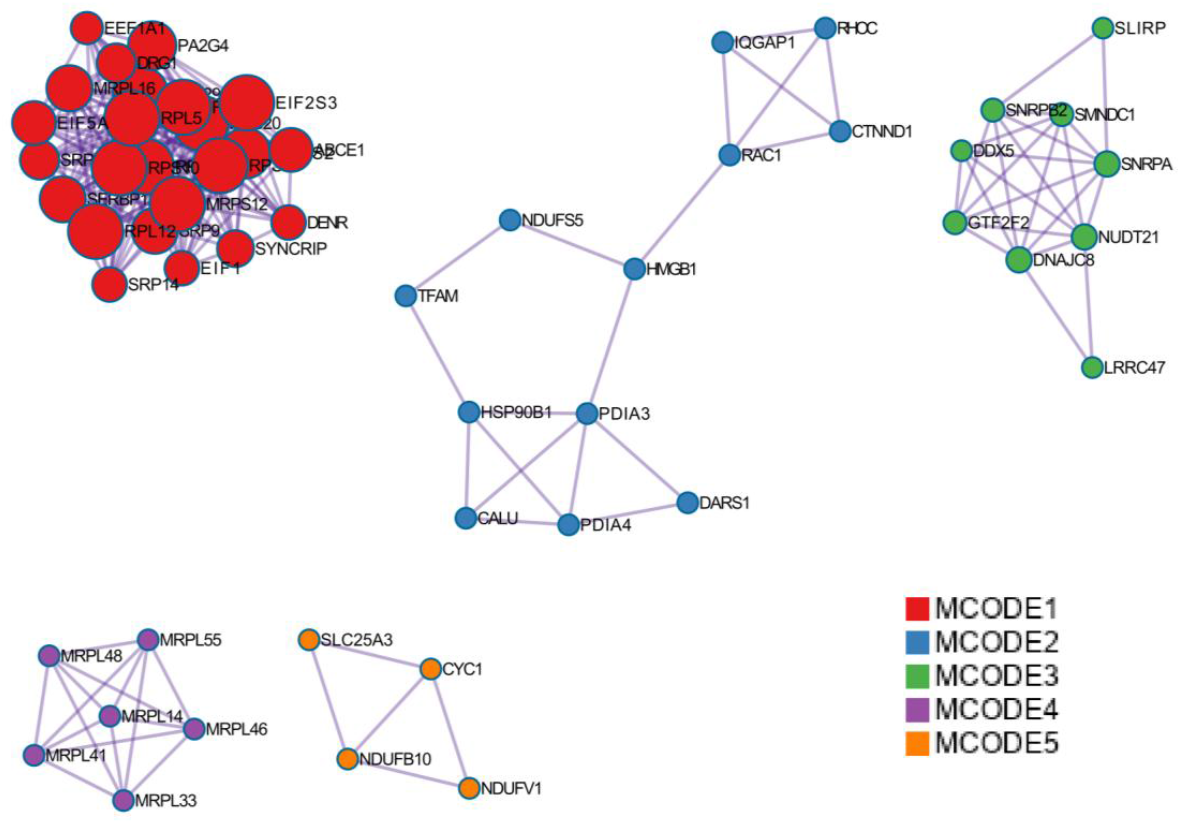
Metascape Molecular Complex Detection (MCODE) subnetworks identified in dataset of 149 proteins enriched more than 40%. MCODE clusters are decoded in Table 1. Detailed results are presented in Supplement 2.

Top 5 putative interactors (Table 2), with highest folds, include MAM-domain containing glycosylphosphatidylinositol anchor protein 1, T-complex protein 1 subunit beta, lysine-tRNA ligase, calumenin and the ribosome biogenesis factor SLX9 homolog.

**Table 2.**
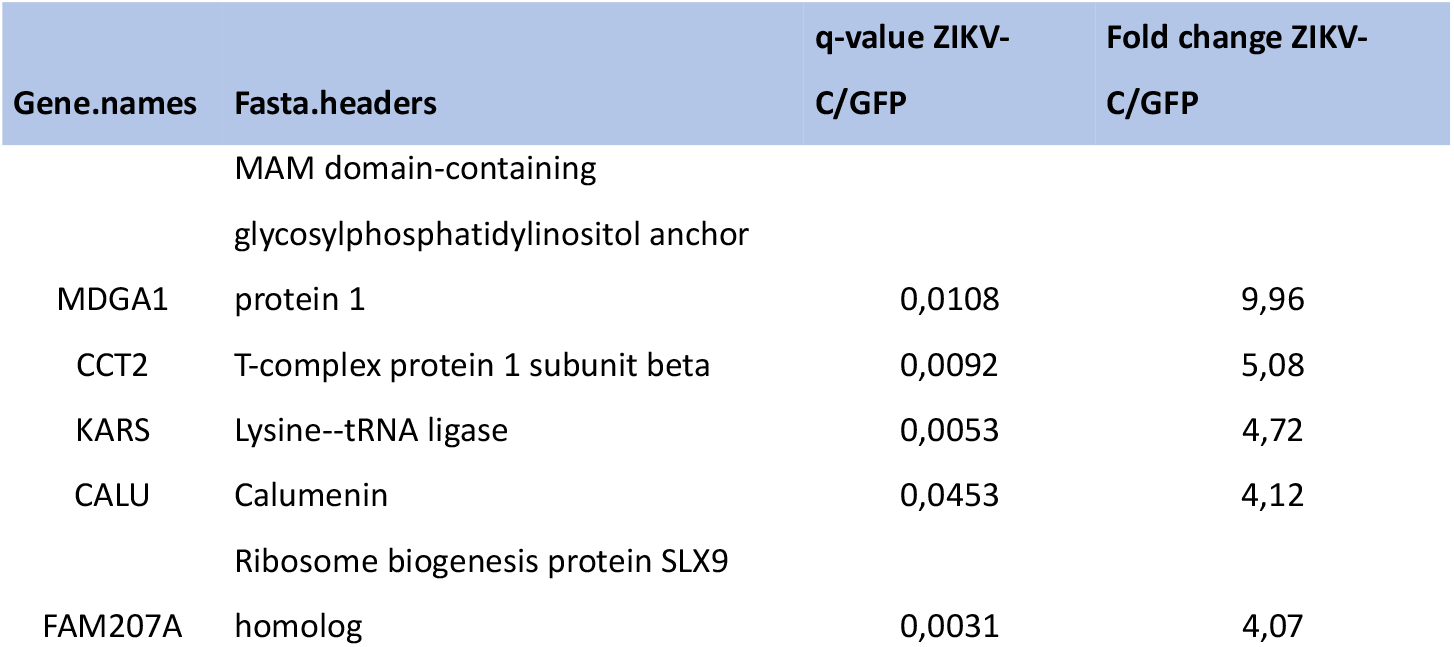
Top 5 putative interactors, with fold changes above 4.

DisGeNet (https://www.disgenet.com/) (Figure 5) analysis of disease-related genes revealed microcephaly and mental retardation (both traits are observed in children born from ZIKV-infected pregnancy) link, associated with eukaryotic translation initiation factor 2 complex. All subunits of this complex are present in putative interactors dataset.

**Figure 5:**
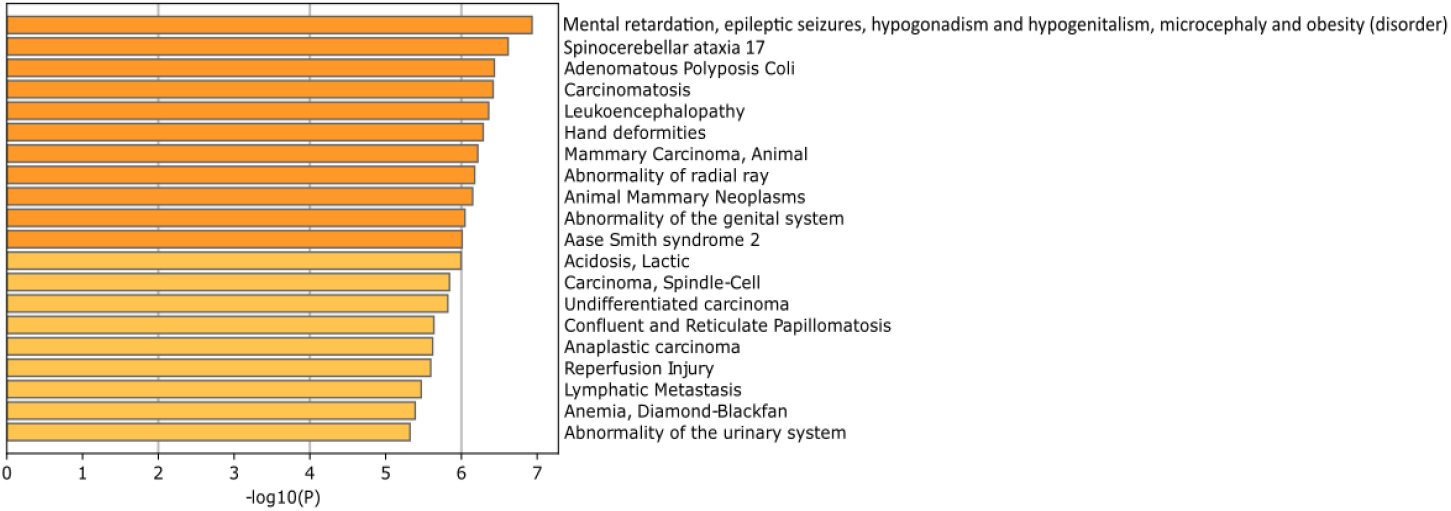
DisGeNet (gene-disease association network) analysis results. The most prominent result is associated with microcephaly, a treat observed in infants born to mothers infected with ZIKA virus.

## Discussion

One of the most important questions with pathogenic viruses such as ZIKA is which elements of the host cell are utilized to ensure the efficiency of the infection cycle. On the other hand, it is also crucial to understand how taking control of the host cells leads to disease symptoms, such as microcephaly in the case of ZIKA. Therefore, protein-protein interaction (PPI) networks of individual proteins of the ZIKA virus and the proteome of mammalian cells, especially in neural stem cells, have been investigated in recent years. A major challenge, however, is the limited overlap among proteomic and functional datasets reported by different research groups. Nevertheless, each of these studies continues to reveal novel molecular mechanisms that may be essential for viral replication and simultaneously detrimental to host cell physiology.. In this study, we present the results of PPI analysis for ZIKA protein C in human neural stem cells. Our findings indicate that the nucleus—particularly the nucleolus—and the mitochondria are major sites of ZIKV-C interaction with host proteins. In turn, control of ribosome biogenesis and protein translation as well as mitochondrial function appear to be the main targets of protein C in the ZIKA virus life cycle in our experimental setup. Furthermore, several identified interactors have known or potential links to neurological and neurodevelopmental disorders. Notably, we report—for the first time—robust interactions between ZIKV-C and MAM domain-containing glycosylphosphatidylinositol anchor protein 1, T-complex protein 1 subunit beta, lysine–tRNA ligase, and calumenin.

Several datasets on the ZIKV protein C interactome in different types of host cells have recently been published^6–9^. However, overlap between these datasets remains low, typically less than 15% in pairwise comparisons. A similar trend was observed in our study, with only 11 proteins shared with the dataset reported by Scaturro et al. [22] (MRPL16, MRPL41, MRPL55, RPL12, RPL5, RPS10, RPS19, RPS20, RPS28, RPS29, RPS5)^22^. These shared interactors are enriched in GO Biological process terms “translation” or “cytoplasmic translation”. On the other hand, 7 and 5 proteins were common to the interactome of ZIKV-C strain variants H/FP/2013 and MR766, respectively^20^. These were MRPL55, FAM207A, DNAJC9, MORF4L2, MRPL48, EIF2A and MRPL16 for strain French Polynesia 2013 H/FP/2013 and MRPL55, FAM207A, MRPL48, EIF2A, WIBG for strain Uganda 1947 MR766. Interestingly, these limited overlaps included mitochondrial ribosomal proteins (MRPL55, MRPL48, MRPL16) suggesting ZIKV-C potential to influence mitochondrial translation. Finally, a comparison of our data with the dataset of Zeng et al.^21^ revealed nine common hits (BUB3, FAM207A, TFAM, SRP14, SNRPA, EIF2S1, SRP9, EIF2S2, HNRNPA1L2).

Although overlap at the individual protein level remains limited, convergence is more apparent when examining the biological processes enriched in these interactomes.. In our study, the predominant categories included regulation of translation (cytoplasmic and mitochondrial), RNA metabolism (processing and splicing), ribosome assembly, rRNA metabolism, and protein targeting to membranes—particularly to the endoplasmic reticulum (ER). Since ribosomal biogenesis occurs in the nucleolus, GO Cellular Component enrichment also included terms associated with this subnuclear structure. These interactions are consistent with previously reported nucleolar localization of ZIKV-C in both overexpression systems and in the context of viral infection^19^. Similar GO enrichment patterns were also identified in interactome analyzes that were reported for capsid proteins of ZIKV or the related Dengue virus^786^. Thus, our dataset fits well with previous studies indicating that protein C regulates host cell proteome to maximize virus production in the ER while protecting viral RNA from anti-viral response^8,10^. However, cytotoxic consequences of such interactions in sensitive cell types such as neural stem cells and immature neurons may include nucleolar stress^19^.

Interestingly, mitochondrial translation emerges as another target of ZIKV-C. Indeed, elongated mitochondria have been observed in ZIKV-infected hepatocytes, an effect attributed to the non-structural protein NS4B, which enhances respiratory function and dampens the antiviral immune response^10^. Moreover, the ZIKV NS5 protein has been shown to bind and inhibit proteins crucial for mitophagy^29^, indicating that the virus may hijack mitochondrial machinery to support replication. ZIKV-C is likely a contributor to this process, and in some cell types, mitochondrial dysregulation may lead to cytotoxicity.

The top ZIKV-C interactors in the current dataset include MAM domain-containing glycosylphosphatidylinositol anchor protein 1, T-complex protein 1 subunit beta, lysine tRNA ligase, calumenin, and the ribosome biogenesis factor SLX9 homolog. Of these, only SLX9 has been previously reported as a C protein interactor^6^, while the others represent novel findings. MAM domain-containing glycosylphosphatidylinositol anchor protein 1, encoded by *MDGA1*, is a glycosylphosphatidylinositol (GPI)-anchored cell surface glycoprotein that plays a role in cell adhesion. It is linked to autism and is highly expressed in the developing nervous system. In mouse models, it has been shown that loss of *Mdga1* expression, specifically in basal neural progenitor cells, reduced NSC proliferation in the subventricular zone (SVZ) and compromised SVZ integrity^11^. Moreover, like other microcephaly phenotypes, MAM domain-containing glycosylphosphatidylinositol anchor protein 1 deficiency is associated with cortical thinning. These disturbances in neuronal development have been attributed to the interaction of MAM domain-containing glycosylphosphatidylinositol anchor protein 1 with connexin 43, a protein essential for the formation of tight junctions^11^. Thus, the interaction of ZIKV-C with MAM domain-containing glycosylphosphatidylinositol anchor protein 1 may contribute to the microcephaly induced by ZIKV infection. Interestingly, studies in non-neuronal cells show that ZIKV infection leads to the destruction of tight or gap junctions and that this effect is due to the loss of connexin 43^12^. Such changes may facilitate ZIKV spread in the infected tissues^13^. Whether ZIKV-C–MDGA1 interaction plays a direct role in this process warrants further investigation..

T-complex protein 1 subunit beta is a group II chaperonin TRiC/CCT, which is responsible for the correct folding of nearly 10% of cytosolic proteins and has been linked to human diseases, including neurodegenerative disorders^14^. It has also been shown to interact with the ZIKV-NS1 protein, promoting viral replication^15^. TRiC/CCT is essential for dengue virus infection^38^ and may regulate NSC proliferation through folding of WD repeat-containing protein 62, mutations in which cause NSC depletion and microcephaly ^16 17,1819^.

Lysine-tRNA ligase is a component of the translation elongation machinery and *KARS1* mutations lead to progressive microcephaly^20^. As in the case of T-complex protein 1 subunit beta, its interaction with ZIKV-C may reduce enzyme activity, impairing neurogenesis. Calumenin is a calcium-binding protein localized in the ER^45^ and belongs to the CREC (Cab45, reticulocalbin, ERC-55 and calumenin) family of proteins. Its cellular functions are closely related to those of the ER and include protein folding and sorting as well as calcium storage and release. Calumenin is an ER stress chaperone that contributes to ER shape and mobility^46–48^. Murine *Calu* gene is highly expressed in the developing brain neurogenic suggesting a role in neurogenesis ^52^. However, future studies are needed to test whether ZIKV-C targeting of T-complex protein 1 subunit beta, Lysine-tRNA ligase, and/or calumenin plays a role in pathogenesis of ZIKV-induced microcephaly.

Finally, several limitations of this study should be acknowledged. First, only one NSC cell line has been used representing one individual human genome. Therefore, natural genetic variation may underlie the apparent variability of ZIKV-C interactions with host cell proteome. Importantly, such a variability may play a role in differential individual sensitivity to ZIKV-mediated neurodevelopmental disruption. Second, the current interactome was established using overexpressed ZIKV-C. However, in ZIKV-infected cells, abundant presence of other components of the virus including its RNA genome and proteins may influence capsid binding to host cell proteome. Likewise, activation of the innate anti-viral response may have additional modulatory effects on ZIKV-C interactions. Therefore, Future studies will be necessary to validate these interactions in the context of natural ZIKV infection and to explore their potential role in pathogenesis..

## Conclusions

We have generated a new dataset for the ZIKV-C interactome in NSC and identified several novel interacting proteins. These findings generate new hypotheses about ZIKV-host cell interactions that facilitate viral replication and disrupt neurodevelopment. In particular, MAM domain-containing glycosylphosphatidylinositol anchor protein 1, the most abundant ZIKV-C-bound protein, has well-documented associations with microcephaly. It remains to be tested whether MAM domain-containing glycosylphosphatidylinositol anchor protein 1 binding to ZIKV-C reduces its pro-neurogenic activity and is sufficient to result in microcephaly in the context of ZIKV infection.

## Abbreviations

ZIKV: Zika virus
WNV: West Nile virus
JEV: Japanese encephalitis virus
NSCs: neural stem cells
NPCs: neuroprogenitor cells
ZIKV-C: protein C from Zika virus
WNV-C: protein C from West Nile virus
JEV-C: protein C from Japanese encephalitis viruses
FDR: False Discovery Rate
PPI: protein-protein interaction
SVZ: subventricular zone
CALU: calumenin
KARS: lysine-tRNA ligase

## Acknowledgements

The authors express their gratitude to Magdalena Blazejczyk, Bartek Frycz and Marta Zurawska for their help with immunoprecipitation experiments, as well as Jacek Oledzki for MS measurements supervision. We also thank Alina Zielinska (IIMCB) for technical assistance and support and Angelika Jocek and Katarzyna Orzol (IIMCB) for laboratory management logistics.

## Funding Statement

Research was supported by Polish National Science Centre Opus grant no. 2018/29/B/NZ2/01752.

## Authors Contributions

AM – mass spectrometry data analysis, writing manuscript

MB – MS sample preparation

AB – co-IP protocol optimization

EL – cell cultures, experimental design

MP – co-IP experiments, experiment design, writing manuscript

BS – MS measurements and optimization, writing manuscript

ES – supervision of co-IP experiments, experimental design, writing manuscript

MH – experimental design, project supervision, writing manuscript

MD – experimental design, project coordination and supervision, writing manuscript, securing funds.

JJ – experimental design, project coordination and supervision, writing manuscript

## Conflict of Interest

None of the authors have any financial or non-financial competing interests.

## Data availability

All data generated or analyzed during this study are included in this published article and its supplementary materials. The mass spectrometry proteomics data have been deposited to the ProteomeXchange Consortium via the PRIDE^21^ partner repository with the dataset identifier PXD064412 and 10.6019/PXD064412.

## Ethics approval

The skin biopsy from a healthy volunteer was obtained with informed and written consent and processed anonymously. The study was approved by the Ethics Committee of the Children’s Memorial Health Institute, Warsaw, Poland (decision no. 112/KBE/2013).

** **Supplement 1** – Supplement1_results.xls file. First sheet (FLAG enriched over 1.4) contains 149 enriched proteins with folds above 1.4. Top 5 putative interactors (highest folds) are marked yellow. All enriched proteins, along with folds, q-values and normalized intensities are presented in sheet “FLAG enriched”, whereas entirety of quantitative and qualitative results is presented in sheets “FLAG_GFP quantification, all” and “FLAG_GFP identification”, respectively.

** **Supplement 2** - Supplement2_metascape_result.upreg_over1_4.xlsx 149 genes inserted into analysis along with their association with MCODE clusters.

## Notes

### Competing Interest Statement

The authors have declared no competing interest.

